# Is the Naked mole-rat a domestic animal?

**DOI:** 10.1101/2022.06.26.497645

**Authors:** Guillermo Serrano Nájera, Koryu Kin

## Abstract

The Naked mole-rat (NMR) is becoming a prominent model organism due to its peculiar traits, such as eusociality, extreme longevity, cancer resistance, and reduced pain sensitivity. It belongs to the African mole-rats (AMRs), a family of subterranean rodents that includes solitary, cooperative breeding and eusocial species. We identified and quantified the domestication syndrome (DS) across AMRs, a set of morphological and behavioural traits significantly more common and pronounced among domesticated animals than in their wild counterparts. Surprisingly, the NMR shows apparent DS traits compared to the solitary AMR. We argue that many of the NMR unconventional traits can be a side-effect of self-domestication. Animals can self-domesticate when a reduction of the fear response is naturally selected, such as in islands with no predators, or to improve the group’s harmony in cooperative breeding species. We propose that self-domestication was necessary to increase social tolerance during the evolution of cooperative breeding and eusociality among AMRs. Finally, we discuss how the DS traits are neutral or beneficial for the subterranean niche and how the increased social tolerance of self-domesticated species could be a side effect of the physical properties of the soil. Our hypothesis provides a novel avenue to enhance the understanding of the extraordinary biology of the NMR.

## Introduction

The reduction of the fight-or-flight response is a hallmark of domestication (1). From the Neolithic period, many animals adapted to the anthropogenic environment by reducing reactivity to humans and perhaps increasing their social tolerance (2). Surprisingly, there is a set of physical traits that often appear in domestic mammals but not in their wild equivalents, known as the domestication syndrome (DS). The DS includes depigmentation, a shorter snout, decreased brain size, floppy ears, reduced or missing teeth and hairlessness, among others traits (3; 4). These traits rarely occur in the wild, and usually, only a subset of them is present in each individual or variety of a domestic species. DS most probably was not directly selected by humans (3); but instead, it appeared as a side effect of selecting for tameness (1). Experiments with wild foxes, rats and minks show that DS traits can appear as a side effect of artificially selecting for tameness (5; 6; 7; 8). However, they can also appear in wild animals when selective pressures favour a reduced fight-or-flight response (self-domestication), as it occurs in islands with no predators (4; 9; 10). Finally, these traits can be observed in primates, including humans (11; 12; 13), bonobos (14), and marmoset monkeys (15), where it has been suggested that the reduced aggression associated with the DS could have been selected to enhance cooperative breeding.

There have been many attempts to explain why selecting for tameness result in the distinct morphological traits encompassed by the DS, including changes in thyroid function (16; 17; 18), in the metabolism of the adrenaline (19; 20; 21), or the retention of juvenile traits in the adult animal (22; 23). Recently, a new, more comprehensive theory proposed that the DS syndrome is derived from mutations affecting the development of the hypothalamic-pituitary-adrenal (HPA) axis and the sympathetic nervous system (24). The HPA axis converts the perception of dangerous stimuli into a hormonal response by transmitting information to the adrenal medulla, which results in the release of glucocorticoids and adrenaline, leading to a long-term elevated reactivity (25). Similarly, the sympathetic nervous system mediates the short-term fight-or-flight response and directly stimulates the adrenal glands. This way, mutations producing a hypofunctional HPA axis could result in the reduced fear response characteristic of domestic animals. Remarkably, the sympathetic nervous system, the adrenal glands, skin melanocytes, bone and cartilage in the skull and teeth, among other tissues, are derived from the neural crest cells (NCC). This way, mild neurocristopathies (disorders derived from defects in the NCC) could explain tameness and the DS simultaneously (24).

The family of African mole-rats (AMR, family *Bathyergidae*) is a well-recognised model to study the evolution of cooperative breeding (26). This family of subterranean rodents contains solitary, social cooperative breeders and two eusocial species: The naked molerat (NMR) and the Damaraland mole-rat. Eusociality is an extreme form of cooperative breeding where species show sexual suppression, reproductive division of labour, overlapping generations and cooperative care of the young (27). In the case of eusocial AMRs, most members renounce to reproduce and specialise in digging, defence, or pup-caring, while a single queen continuously produces offspring (28). The NMR presents the most extreme and unusual traits of its family: it is eusocial, forms extensive colonies (29), and exhibits resistance to hypoxia, delayed senescence, cancer resistance, poor control of body temperature and reduced sensitivity to pain (reviewed in (30)). Its scientific name *Heterocephalus glaber* (“atypical bald head”), also refers to its rare morphological traits: it is hairless (except for sensory hairs), has wrinkled skin, and a flat skull with a reduced number of molars when compared with the other AMR (31). Strikingly, many of these morphological traits can be accounted as part of the DS. Is the NMR a domestic animal?

Here we propose that the social behaviour of AMR is the product of self-domestication. We document that DS traits are more prominent in social AMR species, especially in the NMR. We focus on the NMR because it presents the most extreme morphological and behavioural phenotype, and it is the best-studied member of the family. We propose that many of the unusual NMR traits could be the result of a mild-neurocristopathy that recapitulated the DS. We discuss the possible interactions between the DS and the subterranean niche and argue that eusociality evolved through self-domestication in AMRs. Finally, we argue that this integrative theory will aid NMR research on many fronts.

## Domestication syndrome in the naked mole-rat

The AMR family comprises six genera and more than 15 species. All family members are subterranean rodents and present a complete set of morphological, physiological and behavioural adaptations to the underground environment. However, genera differ in some essential traits regarding their head, digging behaviours, and sociality. Surprisingly, many differences between the social AMR, especially the NMR, and the solitary species can be accounted for as part of the DS. We compiled a list of the DS traits across the AMR family.

### Skull

The skulls of domestic animals presents many characteristic traits. Many domestic animals (e.g. dog, fox, pig, sheep, goat, cat, mouse, cow, human) present a reduced snout in comparison to their wild counterparts (reviewed in (4; 12)). Among subterranean rodents, the relative length of the snout is largely determined by their digging behaviours: forelimb diggers use their claws to scratch soft, sandy soil and possess more prominent faces, while chisel-tooth diggers use their incisors to exploit harder soil and have a relatively wider cranium (32). Chisel-tooth diggers show enlarged zygomatic arcs (cheek bones) and shorter rostra, associated with larger masseter and temporalis muscles, presumably to enhance the bite force (33; 32).

We gathered the width and length of the skull of the AMR, other African Hystricognath (the closest relatives to the AMR: Old World porcupines, the dassie rat and the cane rat) and examples of genetically unrelated convergent mole-rats (from genus *Spalax, Spalacopus* and *Tachyoryctes*). We calculated the ratio between the length and the width of the skull for the chisel-tooth diggers in our data set (Fig. 1, see Materials and methods in the SM). In comparison, exclusive forelimb diggers (other African Hystricognath and *Bathyergus* mole-rats) tend to have a more prominent snout (Supplementary Fig. 1). At the same time, *Tachyoryctes* members, which dig with both their limbs and teeth, present closer cranial proportions to exclusive chisel-tooth diggers (most AMR, *Spalax* and *Spalacopus*). We show that the NMR has a disproportionally shorter snout even for a chisel tooth digger, probably due to a statistically significant reduction of the rostral length (33). The relative shortening of the snout compared to other chisel-tooth diggers is a typical DS trait.

**Fig. 1.**
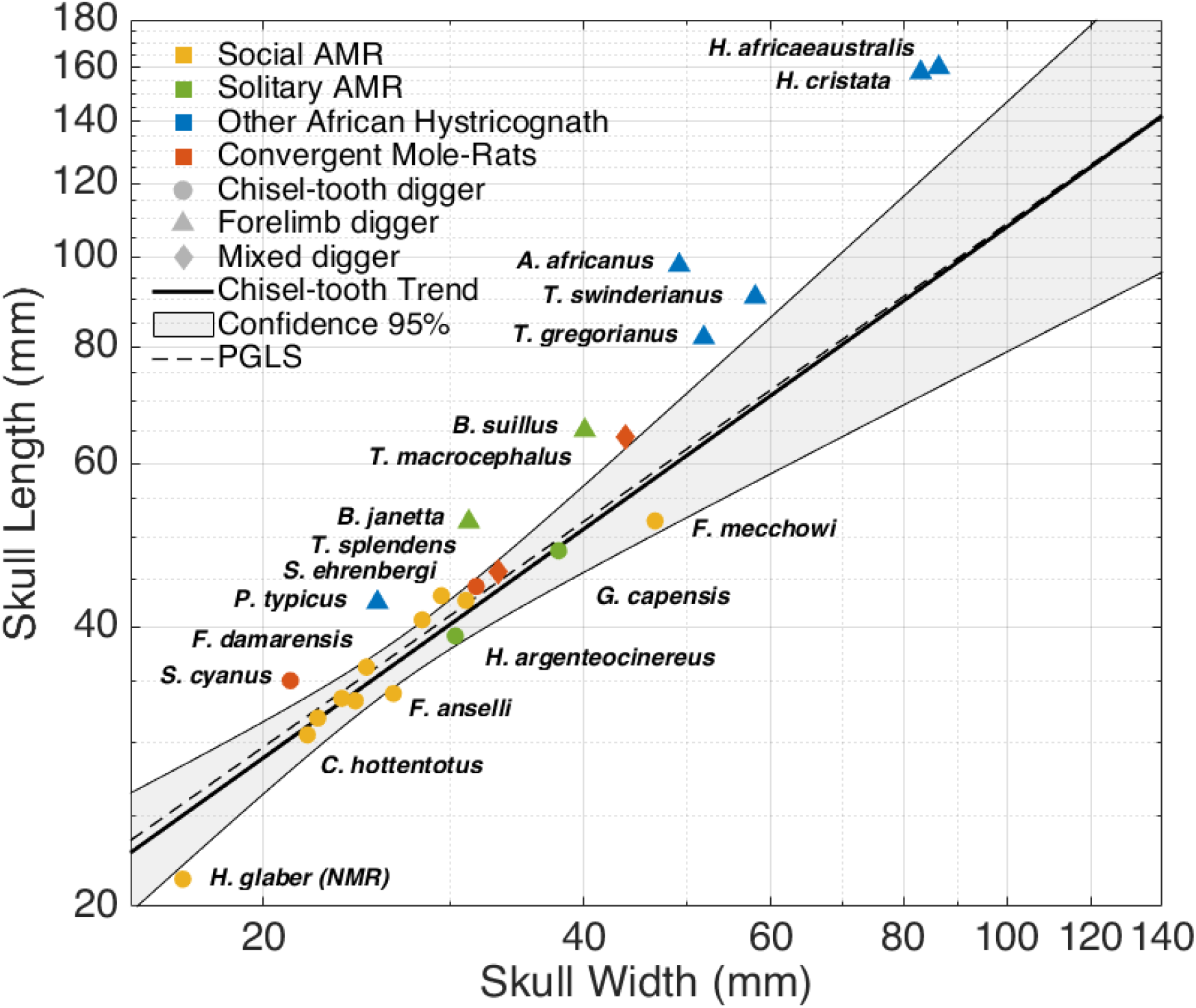
Relation between the length and width of the skull in African mole rats, their closes relatives, and convergent subterranean rodents. Chisel-tooth digging subterranean rodents present a smaller skull length/width ratio than forelimb diggers. The naked mole-rat (*H. glaber*, NMR) has a significantly shorter snout even when compared to other chisel-tooth digging African mole-rats. Each data point is colored according to its social or phylogenetic attributes as indicated in the figure. The shape represents digging mode. The linear regression line for chisel-tooth diggers with 95% confidence intervals is shown, together with the regression line based on phylogenetic generalized least squares (PGLS). All data was retrieved from (34). See Materials and methods in the SM for more details. PGLS was calculated using the evolutionary tree in Supplementary Fig. 2.

All AMR species share a similar diet of geofites, tubers and rhyzomes (34). Unfortunately, there is no data available about the relative hardness of the plants (35) consumed by AMR in comparison with the hardness of the soil. However, it seems unlikely that any tuber or rhyzome offer more resistance than the soil. In addition, while all AMR species eat similar plants (34), they inhabit areas with different soil properties (36). Solitary forelimb diggers (*Bathiergus*), are restricted to more humid environments where digging is easier, while chisel tooth diggers -specially the social species-tend to inhabit regions with harder soils (36). Therefore, it seems that the type of soil correlates better with the head morphology, digging behaviour and sociality (see The subterranean niche and self-domestication), than the type of diet, which suggests that the properties of the soil are the main determinant of the skull proportions among AMR.

### Dentition

The skull of domestic animals often manifests dental abnormalities such as crowding of the cheek teeth or a reduction of teeth size or number (dogs, pigs, mouse, human; reviewed in (4; 12; 37; 38). Most AMR species share the same dental formula (I 1/1, C 0/0, P 1/1, M 3/3; 20 teeth; Table 1), except for *Heliophobus* (39), that shed their premolar teeth and substitutes them with new molars giving a higher, variable number from 20 to 28 teeth; and the NMR (39) which only presents 3 molars (occasionally only 2) in each ramus for a total of 14-16 teeth (34). Therefore, the NMR also presents a reduced, variable number of teeth, as seen in domestic animals.

**Table 1:**
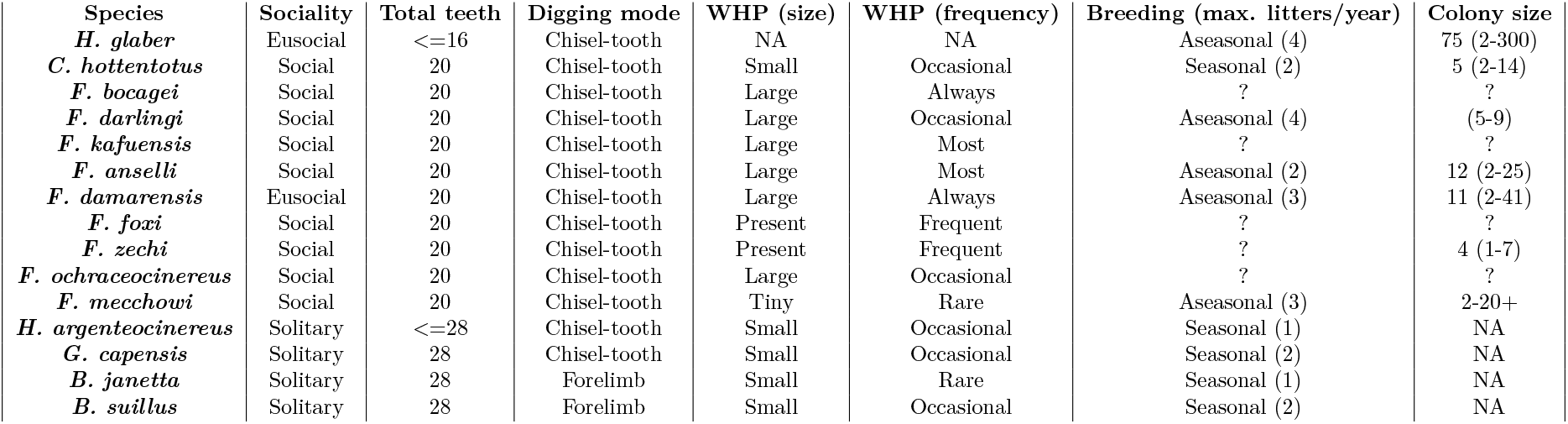
Selected Domestication Syndrome traits in the African Mole Rats. WHP: White forehead patch, not applicable (NA) for NMR because of its complete depigmentation. Colony size is NA for the solitary species. Interrogation mark (?) denotes missing data. All data were retrieved from (34).

### Ear

Another common trait of the DS are reduced or malformed ears (e.g. dogs, horses, pigs, rabbits, cattle, guinea pig; reviewed in (4; 38). NCC contribute to the formation of all the components of the ear, and hearing defects are common in Human neurocristopathies (40). Ears of AMR lack pinna while their closest relatives, the other African Hystricognath, all have well developed external auricles (41). The ear of the NMR appears to be further degenerated compared to other AMRs. They have a very narrow, semi occluded external canal, a smaller than expected and fused malleoincus, a poorly ossified part of the malleus, weakly developed long process of the incus, lack of structural stiffness of the stapes, reduced number of cochlear turns, and a relatively small auditory bulla (42). Furthermore, the ear morphology in the NMR shows a considerable intraspecific variation which could suggest that the ear is subjected to a relaxed selective pressure (42). Finally, some studies indicate that the NMR has relatively poor hearing, while other studies suggest that the auditory system is just adapted to the subterranean niche (reviewed in (30)).

### Brain

Domestic animals (e.g. dog, cat, pig, alpaca, rabbit, mouse, horse) usually have smaller brains than their wild counterparts (reviewed in (4)). Specifically, they tend to show an exacerbated reduction of the forebrain (mink, horse, pig, fox; (43; 24; 44). The brain/body weight ratio is significantly smaller in the NMR when compared to mice (45). Furthermore, the brain size of the AMR has been extensively analysed under the hypothesis that social animals evolve larger brains in (46). However, on the contrary, they found that social AMR species have relatively smaller brains than solitary ones, with a statistically significant reduction in forebrain neurons. In particular, the NMR has a significantly smaller brain mass and fewer neurons in comparison with the other AMR (Figure S1 in (46)). This apparent contradiction could be explained by the self-domestication hypothesis, which instead predicts the smaller brain size of NMR.

### Body and heart

A body size reduction occurred in early domestic breeds (pig, dog, cow, goat, sheep) as a natural adaptation to the human environment (2). It could also be associated with self-domestication in islands (reviewed in (4)), probably as a consequence of small or absent predation and low interspecies competition. The body size of AMR varies exponentially among species. Social AMR species tend to be smaller than the solitary ones, with the NMR being the smallest (~ 30 g) and *B. suillus* the heaviest (~ 1 Kg). In addition to its arguable dwarfism, the body of the NMR differs because it possesses a long tail (50% head-body length), while the other AMR present very short tails (8% – 24% head-body length, depending on the genus; (41)). Similarly, changes in the number of caudal vertebrae occurred in a few domesticated species (dog, cat, sheep; (4)). Moreover, many domestic animals show a shortening of the limbs (4) and, likewise, the legs of the AMRs are generally described as “short and slender” (41), although species-specific measurements are lacking in the literature. Finally, allometric analysis suggests that the NMR has a disproportionally small heart in comparison with mouse (47), a common trait among domesticated species (e.g. mouse, rat, guinea pig, pig, dog; (4)).

### Skin and fur

Probably, complete or partial depigmentation is, along with increased tameness, the most frequent trait in the DS, appearing in at least some varieties of each known domestic species (4). In particular, a white forelock is a common characteristic of domestic animals and a noticeable feature of the Waardenburg syndrome, a human neurocristopathy (24). Furthermore, the white forelock has been recently associated with the self-domestication phenotype in marmoset monkeys (15). Interestingly, most AMR species occasionally present a white forehead patch of variable size (Fig. 2). To our knowledge, the frequency and size of the white patch have not been quantified. However, the available reports (34) suggest that the white patch tends to be more prominent and frequent among social species and reduced and only occasionally present in the solitary mole-rats (Table 1). The NMR appearance departs from the other AMRs because of its complete depigmentation, hairlessness, and skin folds. However, all these attributes are also characteristic of some domestic variates (dog, cow, pig, goat, rabbit; (4)) and can be understood as a more extreme domestication pheno-type.

**Fig. 2.**
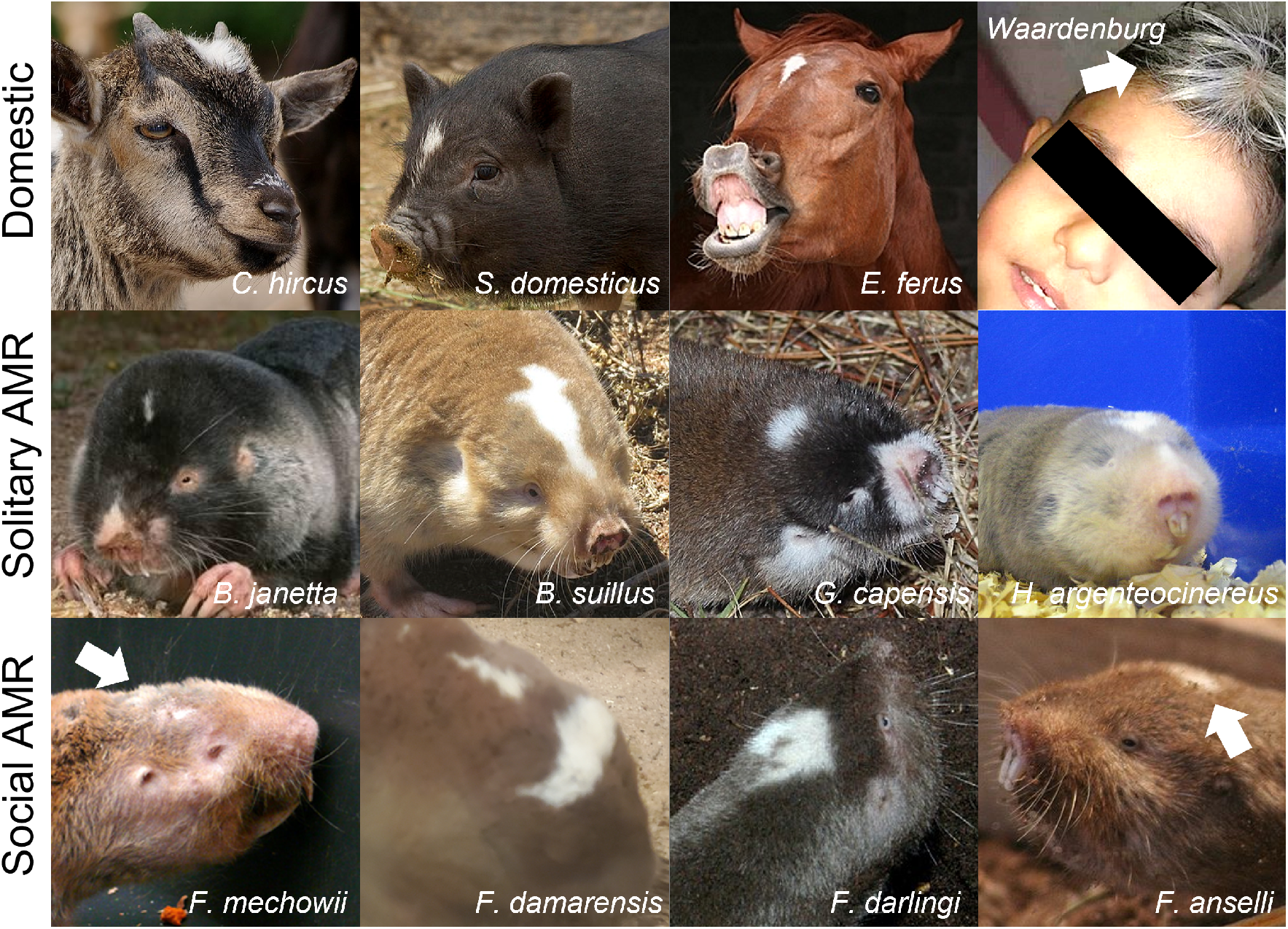
White forehead patch in domestic species and African mole rats. Domestic species and some neurocristopathies are associated with large white patches on the forehead. Most African mole-rats present a white forehead patch which size and frequency could be associated with their social behaviour (Table 1). All images are available under a CC BY. Authorship credit of the images in Supplementary Table 1.

### Reproductive biology

A combination of body size reduction, earlier sexual maturation, increased litter size, non-seasonal oestrus cycles, and a decrease in sexual dimorphism generally allow domestic animals to have higher reproductive rates than their wild counterparts (2; 4). Non-seasonal reproduction, higher annual reproductive output, and absent or reduced sexual dimorphism seem to be more frequent among social AMR (*Cryptomys, Fukomys, Heterocephalus*). The lack of behavioural and morphological sexual dimorphism is exceptionally noticeable in the non-reproductive members of a NMR colony (48): The males lack scrotum, the females have an imperforated vagina; and, unlike any other mammal, the phallus (clitoris or penis), the anogenital distance, and the perineal muscles are sexually monomorphic (29; 49; 50).

They also lack sexual dimorphism in femoral bone structure and quality (51) and forebrain regions (52), which has been associated with the retention of a pre-pubertal phenotype in the worker castes (53; 54; 55).

While domestic animals seem to present an earlier sexual maturity, most of the individuals in a NMR colony never reproduce, and they remain all their life sexually suppressed in a pre-pubertal state until the breeding members need to be replaced (56; 57). This phenomenon can be seen as an extreme version of neoteny. Indeed, neoteny has been classically used to explain many of the characteristic reproductive and behavioural changes that occur during domestication (4). Many of the characteristic traits of the NMR can be considered neotenic (58) such as the lack of hair, absence of scrotum or the decline of bone mineralisation with age (59), which could be simultaneously related to the extraordinary social organisation and longevity of this species.

### Hormones and social behaviour

Glucocorticoids mediate the long-term response to environmental and social stressful situations. A reduction of the concentration of corticosteroid hormones characterises both domestic animals (1; 5; 24; 60) and subordinate individuals in cooperative breeding species (61; 25). In domesticated animals, the reduction of glucocorticoids could lower aggressive behaviours and increase social tolerance, which could also be beneficial to maintain harmony within the group in cooperative breeding species. In this line, both eusocial species *F. Damarensis* (62) and the NMR show little aggression among individuals of the same colony (63). However, while *F. Damarensis* manifest lower aggression against unfamiliar conspecifics, the NMR shows xenophobic behaviour (63; 64; 65).

In NMR colonies, the queen displays most aggressive behaviours (66); however, cortisol levels, the main glucocorticoid in the non-murine rodents (67), do not correlate with social or reproductive status either in the NMR (68; 66) or *F. damarensis* (69) which suggest that social AMR do not suffer “stress of dominance” or “stress of subordination” (68).

Nonetheless, cortisol levels rise after the death of the queen producing a violent period when individuals fight for the succession (70; 71), or in an individual isolated from the rest of the colony (72; 68), indicating that they suffer from social stress as observed in marmoset monkeys (73) and tamarins (74), two cooperative breeding primates. Finally, in AMR species, there are no significant differences in the basal blood concentration of cortisol among non-pregnant females in one solitary species (*G. capensis*) and three social species (*F. darlingi, F. hottentotus pretoriae, F. damarensis*). However, only the solitary species showed an elevation of cortisol levels after repeated encounters with unfamiliar conspecific of the same sex (62). Altogether, these observations suggests that cooperative breeding species suppressed social stress in order to acquire social tolerance towards members of the same species.

Oxytocin is also a mediator of the HPA axis that has been associated with increased social cohesion and maternal behaviour in domesticated animals (60). In the same line, the NMR expresses more oxytocin receptor levels in the nucleus accumbens, a forebrain centre controlling monogamic, maternal and allomaternal behaviour in comparison with the solitary *G. capensis* (75). Moreover, non-breeding individuals of a NMR colony present more oxytocin-producing neurons than breeding individuals, suggesting that oxytocin contributes to maintaining the prosocial behaviour of the worker caste (76).

## Other traits linked to an altered NCC functionality

The NMR shows a series of peculiar biological traits that are not generally included in the DS, but that could be explained through defects in the physiology of the NCC.

### Peripheral nervous system

Most of the structures of the peripheral nervous system (cranial and spinal nerves and ganglia; and enteric nervous system) are derived from the NCC (77). The NMR also manifest alterations in the cutaneous innervation. It presents a significantly reduced ratio of unmyelinated nociceptive C-fibres and A-fibres in the saphenous and sural nerves (1.5:1), while in most mammals C-fibres generally are 3-4 times more abundant (78). Innervation defects are typical of certain neurocrystopathies (79) and could be manifestation of mutations affecting the biology of NCC (see Trka in the main text).

### Thymus

NCC participate in the development of the stroma of the thymus, an organ necessary for the maturation of the T-cells (80); and consequently, the T-cell population is affected in several neurocristopathies (81; 82). The NMR has a reduced thymus/body weight ratio and presents a poor delineation between cortex and medulla in comparison to mice (83). Moreover, the immune system of the NMR differs from mice and humans in having predominantly myeloid cells and a notable deficit in cytotoxic T-cells (84), which could be related to a defective thymus development due to alterations in the NCC.

### Thyroid

The NCC also contribute to creating the stroma of the thyroids (85) and altered thyroid function is a common trait among neurocristopathies (81), and it has been previously associated with domestication (16). Interestingly, the NMR exhibits unique low levels of thyroid hormone (86; 67).

### Gall bladder

Gall bladder derives from the NCC (87). Although most vertebrates possess this gland, it is absent in the NMR (88).

## Gene candidates

Here we discuss some well-studied mutations in the NMR that are associated with the NCC physiology or with the control of the fear response.

### Trka

Trka (tropomyosin receptor kinase A) binds to NGF (nerve growth factor), promoting survival and differentiation of the sensory neurons during embryonic development, and modulating the sensitivity to pain later in life (89). Trka is implicated in cell migration in normal and pathological conditions (90); and it is necessary for the formation of the dorsal root and sympathetic ganglia by the trunk NCC in mouse (91).

In humans, mutations in the Trka gene produce congenital insensitivity to pain (CIP) with anhidrosis (CIPA), a neurocristopathy that results in a total loss of the C-fibres, lack of nociception, inability to sweat (hypotrophic sweat glands), lack of hair (bald patches in the scalp), poor thermoregulation, and oral and craniofacial manifestation including missing teeth and nasal malformations (92; 79). Furthermore, CIPA patients seem to be unable to learn new fears and probably fail to exhibit a fight-or-flight response (79).

Knock out mice for Trka present a similar phenotype: Abnormal peripheral small nerves fibres, deficient nociception, small body size, early death, and severe cell loss in trigeminal, dorsal root, and sympathetic ganglia. However, they do not manifest anhidrosis, and over time they develop a mottled fur with scabs (91; 79). Trka null mice also show an acute decrease of the basal forebrain cholinergic neurons (BFCNs). Notably, BFCNs have been implicated in the control of conditioned fear behaviours (93), and mice lacking Trka signalling in the forebrain show defective fear conditioning (94). Furthermore, Trka null mice show a defective thymus formation with no clear delimitations between the cortex and the stroma and a reduced number of thymocites (95).

The NCC hypothesis states that domesticated animals suffer a mild neurocristopathy produced by mutations that only partially reduce the activity of genes implicated in the NCC biology (24). The NMR possess between one and three amino acid substitutions in the kinase domain of Trka that turn the receptor hypofunctional that are absent or rare in other animals or among other AMR (96). Moreover, the expression level of Trka in the NMR is significantly lower than that of mouse (97). It is tempting to see many of the traits of the NMR as partly produced by a mild version of CIPA, caused by the reduced expression of a hypofunctional version of Trka. In this line, PRDM12 (PRDI-BF1 and RIZ homology domain-containing protein 12), a transcription factor downstream of Trka also associated with CIP in humans (98), shows lower expression and unique aminoacid variants in the NMR (97). Altogether, mutations in the Trka pathway could explain the reduction in the C-fibres and the reduction in pain sensitivity in the NMR (96); but also the lack of fur and sweat glands, the poor thermoregulation, the reduced and defective thymus, missing teeth, shorter muzzle and the smaller forebrain. Indeed, Trka could provide a genetic connection between the neural crest and the reduction of the forebrain suggested by the neural crest domestication hypothesis (24). Finally, Trka mutations also seem to cause a reduction in the fear response in mice and humans, and could potentially explain the high social tolerance of the NMR.

### HIF1*α*

The NMR presents critical molecular adaptations to survive in hypoxic/hypercapnic environments produced by large numbers of individuals living in borrows (99). For example, the NMR carries a hyperactive version of HIF1*α* (Hypoxia Inducible factor 1 *α*), a transcription factor that regulates the expression of dozens of genes in response to low-oxygen conditions. In normal-oxygen conditions, HIF1*α* is targeted for degradation by VHL (Von Hippel–Lindau tumor suppressor); however, low-oxygen levels prevent the HIF1 *α*-VHL interaction. The NMR possesses a unique mutation in HIF1*α* (T395I, Fig. 3) that prevents its interaction with VHL, as well as a mutation in VHL (V166I) which probably further reduces the ubiquitination of HIF1*α* (100). Moreover, the NMR expresses higher levels of HIF1*α* than mice in several tissues (101). Lastly, HIF1*α* probably promotes a glycolytic metabolism (102; 103), a less efficient pathway to obtain ATP than oxidative phosphorylation that works in the absence of oxygen. Glycolysis produces lactic acid as a by-product, which blocks glycolysis by inhibiting PFK1 (phosphofructokinase-1). The NMR presents a unique adaptation to bypass the glycolysis block during hypoxia by using fructose (instead of glucose) as a substrate, enabling the glycolysis even in high lactic-acid conditions (104).

**Fig. 3.**
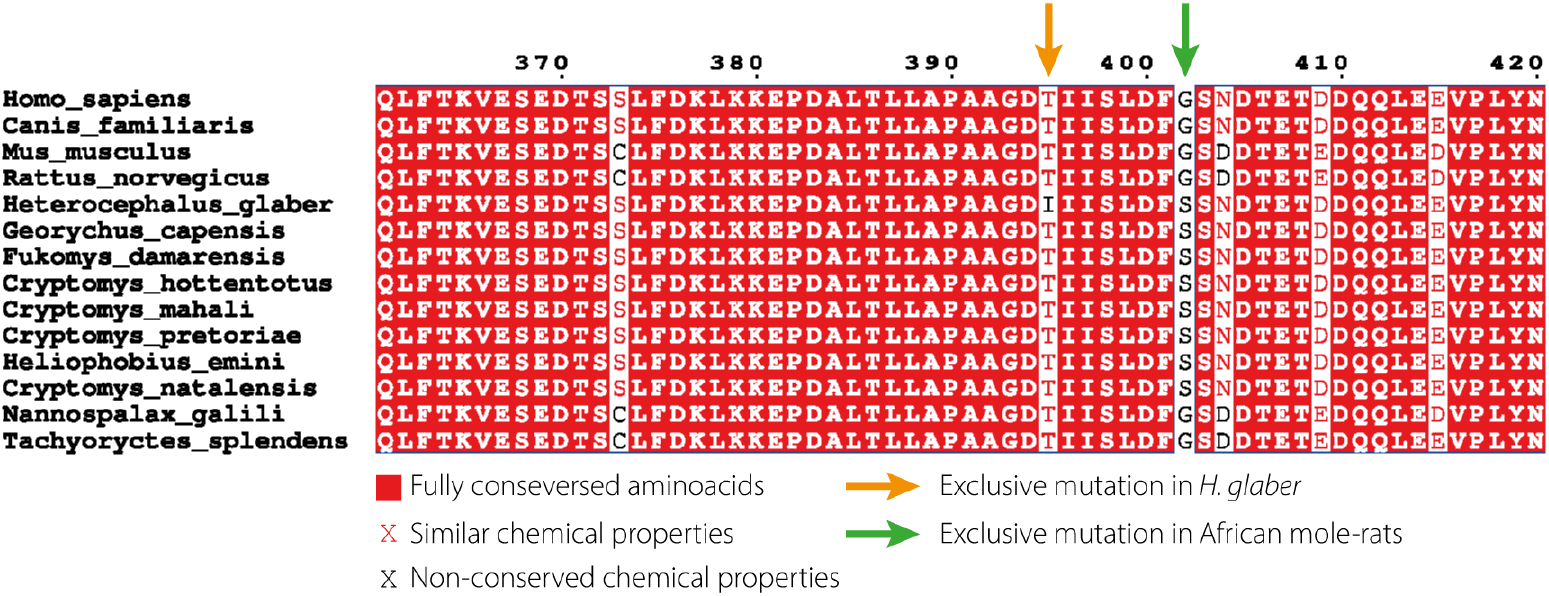
Conservation of HIF1*α* among African mole-rat and other subterranean rodents. The naked mole-rat (*H. glaber*) presents a unique mutation (T395I) that presumably prevents being targeted for degradation. Alignment generated with ENDscript2 (126) using data from (127)

Recently, it has been suggested that metabolic regulation is essential for the migration and differentiation of the NCC (105). Indeed, genetic disorders caused by mutations affecting the glucose metabolism produce severe congenital disabilities with severe craniofacial deformations similar to neurocristopathies (106; 107). NCC switch to glycolytic metabolism during delamination and migration and return to oxidative phosphorylation during differentiation (105). In fact, HIF1*α* upregulates many NCC-specific genes and increased, and reduced HIF1*α* activity results in defects in the NCC migration (108). Furthermore, alterations in VHL are associated with the von Hippel-Lindau syndrome, a neurocristopathy that can cause a variety of neoplasms, including the abnormal tissue growth of the adrenal gland (81). Intriguingly, more than the 20% of the NMRs in a zoo study presented adrenal hyperplasia (109).

### ASIC4

The NMR also presents adaptations to protect the brain against acidification. Acidotoxicity occurs when the accumulation of lactic acid during hypoxia acidify the medium to pathological levels, triggering the *Ca*^2+^ ASICs (acid-sensing ion channels) (110). The NMR shows a significantly reduced expression of ASIC4 in the brain compared to mouse, which protects its brain during acidosis (111). Furthermore, a recent report has identified ASIC4 as a marker of the early differentiation of the NCC (112), and ASICs expression has been implicated in the processing of the fear response (113; 114; 115). In particular, ASIC4 in mice seems to modulate the freezing behaviour in the presence of predator odour, but not the shock-evoked fear learning (116). This way, mutations affecting the expression of ASICs could reduce the fight-or-flight response in the NMR giving rise to a more docile phenotype.

## The subterranean niche and self-domestication

Although AMR and NMR show many traits expected in underground rodents (fusiform body, short limbs, reduced external protuberances (32)), many traits seen in NMR cannot be simply explained by adaptation to subterranean niche (reduction of the forebrain, hairlessness, lack of sexual dimorphism, reduction of pain sensitivity, immature thymus…).

Traditionally, the DS has been explained as a pleiotropic side effect of selecting for tameness (1; 24). However, in some self-domesticated species, it is conceivable that the DS appears as a side effect of selecting for other trait associated with the NCC biology. We propose that harder soils, in conjunction with the subterranean niche, fostered the evolution the DS in AMR. AMR ancestors evolved more solitary and aggressive behaviours -a *wilder-like* phenotype-as they radiated towards more humid areas and encountered more resources and softer soils (36). Indeed, while the solitary AMR (*Bathiergus*, *Georychus*, and *Heliophobus*) are restricted to more humid environments where digging is easier, the social AMR (*Fukomys* and *Cryptomys*) mostly inhabit coarse sandy soils and the eusocial AMR (NMR and Damaraland mole-rat) tend to live in areas with hard, compacted soils (36).

The hardness of soil is often associated with the mode of digging in subterranean rodents. Curiously, among AMR, the only forelimb digger genus (*Bathyergus*) is solitary and territorially aggressive, while all social species (*Fukomys, Cryptomys, Heterocephalus*) are chisel-tooth diggers (34). Chisel-tooth digging is an adaptation to harder soils that requires a wider skull to enhance the attachment of larger muscles (33; 32). Potentially, harder soils could favour the widening of the rostrum, by creating an evolutionary pressure on the cranial NCC that resulted in the appearance of morphological and behavioural side effects associated with the DS. Simultaneously, harder soils can be better exploited in cooperation with other individuals to reduce the cost of digging extensive tunnels and improving the chances of finding dispersed food (36). Furthermore, molecular adaptations to the hypoxic environment produced by large colonies living underground could have an important impact in the physiology of the NCC, producing a more exacerbated DS. Finally, some of the characteristic traits of the DS traits could be either beneficial or neutral for a subterranean animal: the dwarfism, reduction of the ear, shortening of the limbs and flattening of the rostrum could be beneficial for moving through tunnels and digging, while alterations in colouration are probably less critical to animals living underground. Therefore, we argue that the that genes associated with the physiology of the NCC in the NMR have been under evolutionary pressure (Fig. 4).

**Fig. 4.**
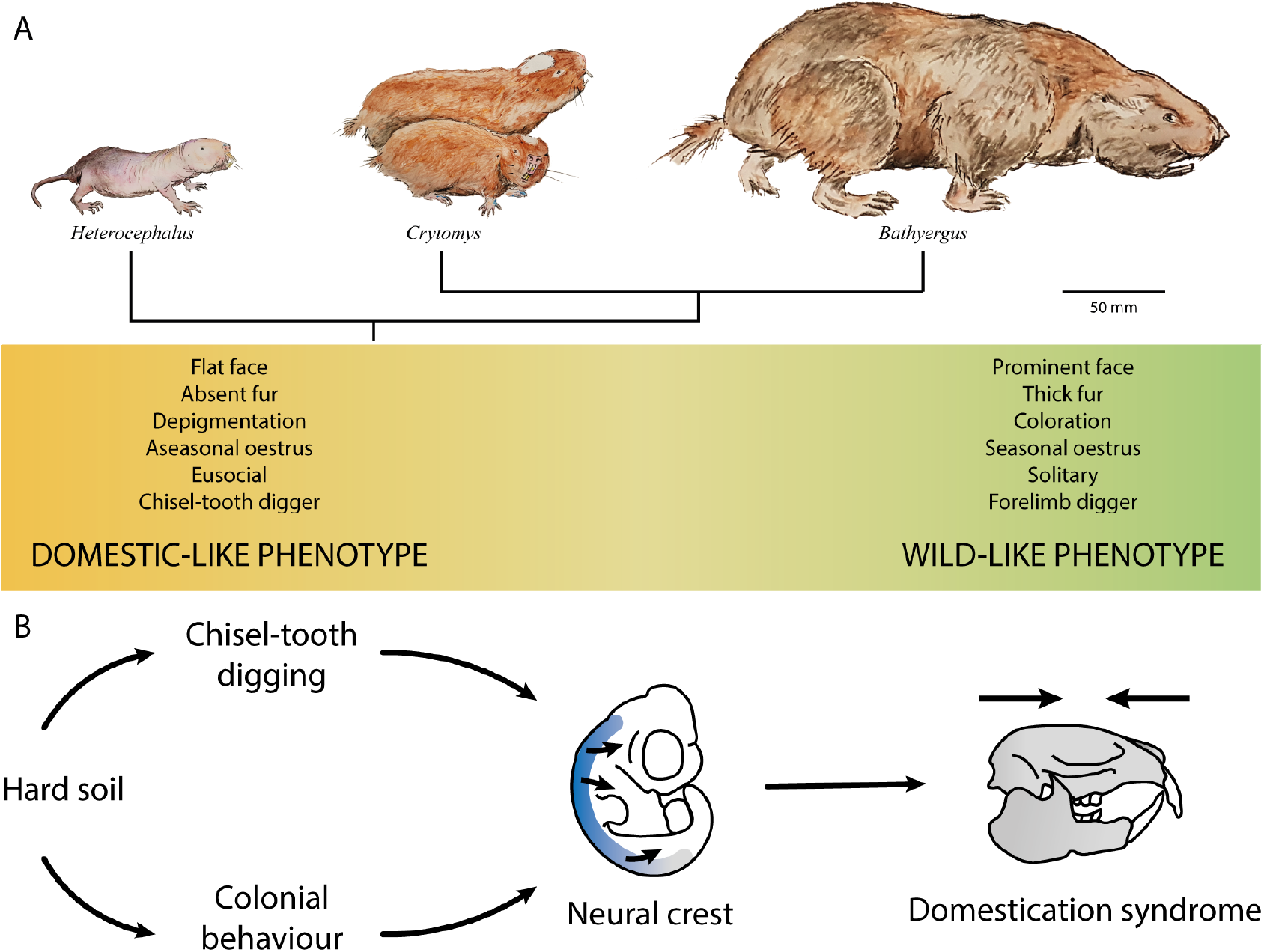
Domestication syndrome in subterranean rodents. A) Evolution of the African mole-rats from a *domestic* to a *wild* morph. B) The subterranean niche could promote self-domestication. Chisel-tooth digging facilitates breaking hard ground, and a colonial, cooperative life-style reduces the amount of work carried out of each individual. Therefore, coloniality and chisel-tooth digging could be adaptations to hard soils. Coloniality needs a high degree of social tolerance, and chisel-tooth digging needs flat faces for large muscle attachments, which could serve as selective pressures foe the neural crest producing the appearance of the domestication syndrome in subterranean rodents. Furthermore, large number of individuals underground generate a hypoxic environment which could favor the appearance of defects in the neural crest development.

Altogether, this suggests that the subterranean niche could promote the self-domestication of underground rodents. In fact, genetically unrelated, convergent subterranean species could also present signs of self-domestication. For example, *Spalax ehrenbergi* is a superspecies of subterranean rodents that inhabit the eastern Mediterranean from Israel to Egypt. Similarly to AMRs, they present morphological adaptations to living underground such as short limbs, wide crania (Fig. 1), with varying coat colouration with occasional white forehead patches (117). Strikingly, although they are described as solitary, chromosomally distinct species present different ratios of aggressive and “pacifist” individuals. The Egyptian Spalax variant, which includes only “pacifist” animals, inhabits the aridest environment (118) and can be distinguished by its shorter mandible (119). Furthermore, they present convergent genetic adaptations to the hypoxic environment (120; 121).

## Testing the NMR self-domestication hypothesis

The self-domestication hypothesis for NMR presented in this paper can be tested in various ways. First, new behavioural studies are necessary to evaluate the fear response in different AMR species, for example through the evaluation of the freezing behaviour in classic fear-conditioning experiments where a mild electric foot-shock is applied after an acoustic stimulus. However, the voltage threshold must be carefully calibrated for each species because they potentially present very different pain sensitivities (97). Alternatively, the freezing behaviour could be measured in the presence of the odour of a predator relevant for AMR species, for example, using snake or small carnivore faeces (34). We expect AMR with a more domesticated phenotype to present an altered fear-conditioned behaviour compared to solitary, forelimb diggers. Furthermore, it is difficult to compare the aggressiveness and fear response of eusocial and solitary species, because cooperative breeders can be affected by their ranking a reproductive status (65), variables that should be controlled. Gathering data on the size and activity of the adrenal glands and stress hormones during the fear response across different species could also be informative to test this hypothesis.

To test the implication of NCC in the evolution of AMRs we propose inter-species GFP-labelled NCC grafting experiments (24). We expect NCC from the NMR to show a deficient migration and/or differentiation in comparison with *Bathyergus* mole-rats. In addition, we expect that phylogenetic analysis of genes undergoing positive and relaxed selection (122) among AMR will highlight genes known for their involvement in NCC development or in neurocristopathies.

Finally, we propose the generation of an extensive database of subterranean and fossorial rodents containing their digging mode and other relevant morphological and behavioural traits. Using this database in combination with statistical analysis corrected by phylogeny to avoid evolutionary relationships affecting the independence of the individual samples (123), it would be possible to test if the digging mode in subterranean rodents is associated with DS traits and with an increased social behaviour.

## Discussion

We showed that the NMR when compared with solitary AMRs, presents a phenotype compatible with the DS. We suggest that the DS could be beneficial for the evolution of life underground; and that the digging mode, determined by the properties of the soil, could create an evolutionary pressure on the NCC that generated the DS among AMRs. Furthermore, we highlighted some known mutations compatible with the evolution of a domestic phenotype and proposed experiments to test our hypothesis.

Taken all these together, we propose that the high degree of social tolerance necessary for the evolution of cooperative breeding and eusociality in AMRs occurred through self-domestication. A combination of scarce resources in increasingly arid environments (36; 26), together with the constraints imposed by hard soils and the underground environment, could create selective pressures that directly and indirectly lead to the self-domestication of the AMR. On one hand, an increased social tolerance associated with self-domestication could be beneficial for subterranean rodents because a colonial, cooperative life-style reduces the digging work carried out for any individual and increases the chances to find disperse food (36). Hard soils could impose an evolutionary pressure toward chisel-tooth digging, which requires a flatter face in order to increase the muscle attachments. If the muzzle reduction occurs through the selection of less active NCC, this could generate the pleiotropic effects seen in domestic animals and in neurocristopathies, and might increase the social tolerance as a side effect. Moreover, the gathering of large number of individuals in subterranean tunnels generates a hypoxic environment which could lead to molecular adaptations that affect to the NCC development, further enhancing a self-domesticated phenotype. Intriguingly, in this way, the properties of the soil could act simultaneously as a distal (36) and a proximal factor for the evolution of sociality. In other words, the evolution of sociality in subterranean rodents could be initiated as a side effect of the underground environment and properties of the soil, providing an explanation for the abundance of colonial life among fossorial species.

Previously, neoteny has been used as a unifying biological explanation of the many distinctive NMR’s morphological and physiological traits (58). Nevertheless, it is unclear what evolutionary pressures guided the evolution of a neotenic phenotype or how neotenic traits relate to each other. Instead, we argue that the self-domestication hypothesis is a more comprehensive framework for the evolution of the NMR. In fact, neoteny is often associated with a domestic phenotype, and neotenic and DS traits can be confounded (4). Our hypothesis offers a clear rationale to explain many NMR’s attributes, how they relate to each other and pinpoints the evolutionary pressures that drove them.

The DS can be controversial due to the lack of a consistent definition of the traits among different publications (38). Under the mild neurocristopathy theory of domestication, the DS is seen as a collection of pleiotropic effects of variable penetrance produced by mutations affecting the NCC, as can be readily seen in human neurocristopathies (24; 81). Furthermore, in viable animals, the effects of the mutations affecting the NCC can be further confounded by compensatory mutations in some of the altered organs (124). For example, all the components of the ear in the NMR show signs of degeneration and intraspecific variation (42), however the NMR accumulated positively selected mutations affecting the cochlear hair bundles (125). A plausible interpretation is that the general ear degeneration produce by a mild-neurocrystopathy was later compensated by mutations that readapted the NMR hearing to the subterranean environment. In addition, some other traits associated with DS are neutral in the subterranean niche, and as such, they will not be compensated. This relaxed selective pressure in the underground environment could produce the exacerbation of certain traits, explaining perhaps the highly variable white forehead patch or the defects in the pain perception in the AMR family. Thus, the AMR could show how the combination of positive compensatory selection and relaxed selection can produce variations within the DS.

The study of the NCC could guide future research in the NMR as an animal model and provide new clues to understand its extraordinary biological traits. The recognition of self-domestication and its pleiotropic effects could lead to a deeper understanding of the interactions of social behaviour and the physical environment during evolution.

## Acknowledgments

GSN acknowledges support from an EASTBIO BBSRC PhD student training grant (1785593). KK acknowledges support through JSPS Overseas Research Fellowship (H28–1002) and la Caixa postdoctoral juniour Leader fellowship (LCF/BQ/PI20/11760009). Both authors thank Kees Weijer for his comments and support, and Juan Pedro Fernández Doctor for the illustrations of the African mole-rats.

## Supplementary material

### Materials and methods

Data in Table 1 and the mean values of skull lengths and widths for each species in Fig. 1 and Supplementary Fig. 1, data in were retrieved from (1). For species with conspicuous sexual dimorphism, only the values for the male organisms were used .

The dataset was analysed with a statistical package R (2). Linear regression based on ordinary least squares and its confidence interval was computed with an R function “lm”. Linear regression based on phylogenetic generalized least squares (PGLS) was performed with an R package “nlme” (3) and “phytools” (4). The tree used for the PGLS analysis is based on the node-dated exponential tree in (5), from which the species not included in this study were pruned away (Supplementary Fig. 2).

## Supplementary Figures

**Supplementary Fig. 1.**
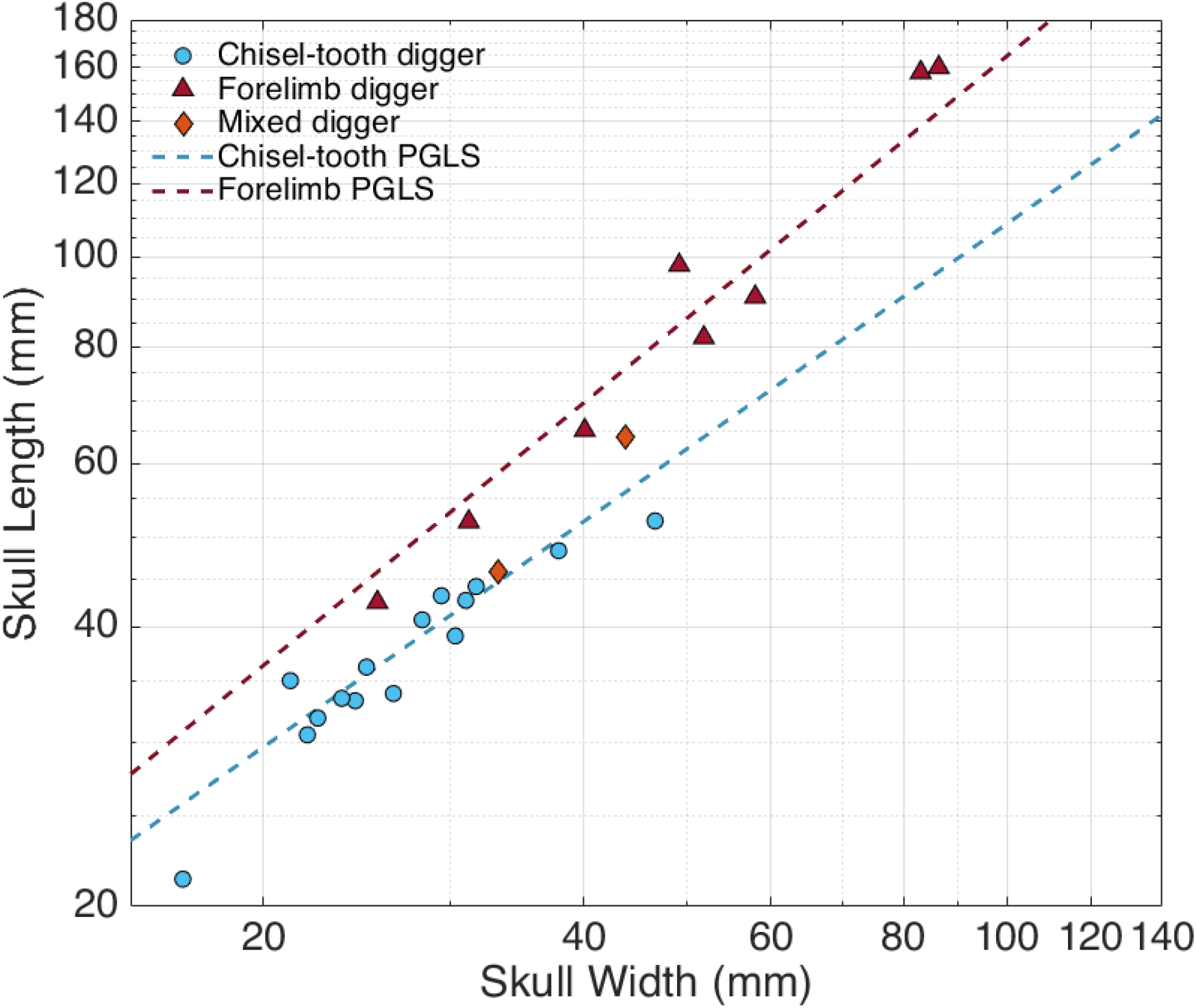
Trends in chisel-tooth and forelimb digger head proportions. Phylogenetic generalized least squares regression (PGLS) of the head proportions in chisel-tooth and forelimb diggers. Chisel-tooth diggers present a flatter face. PGLS was calculated using the evolutionary tree in Supplementary Fig. 2.

**Supplementary Fig. 2.**
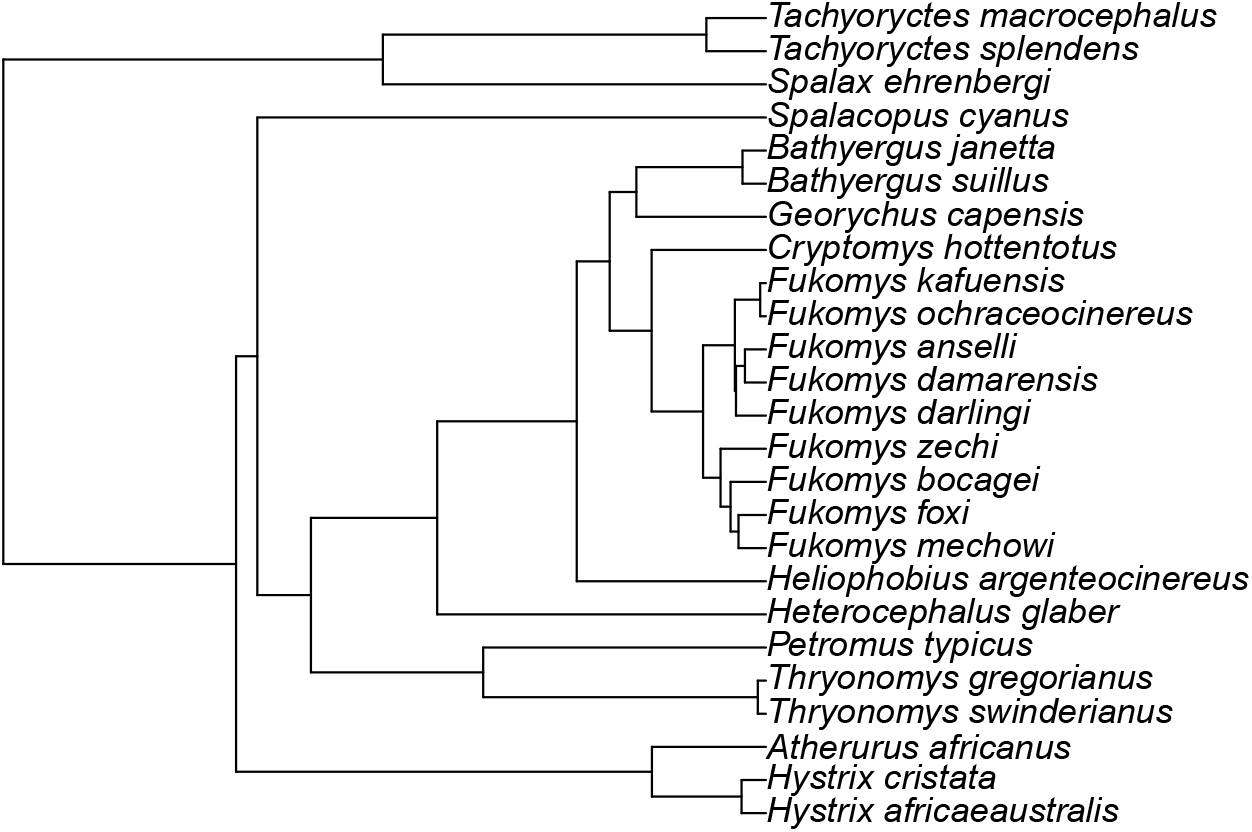
Phylogenetic tree of the AMR, its closes relatives and other convergent mole-rat species. This tree, which is based on the node-dated exponential tree in (3), was used to compute the phylogenetic generalized least squares regression (PGLS) of the head proportions (Fig. 1, PGLS and Supplementary Fig. 1).

**Supplementary Table 1.**
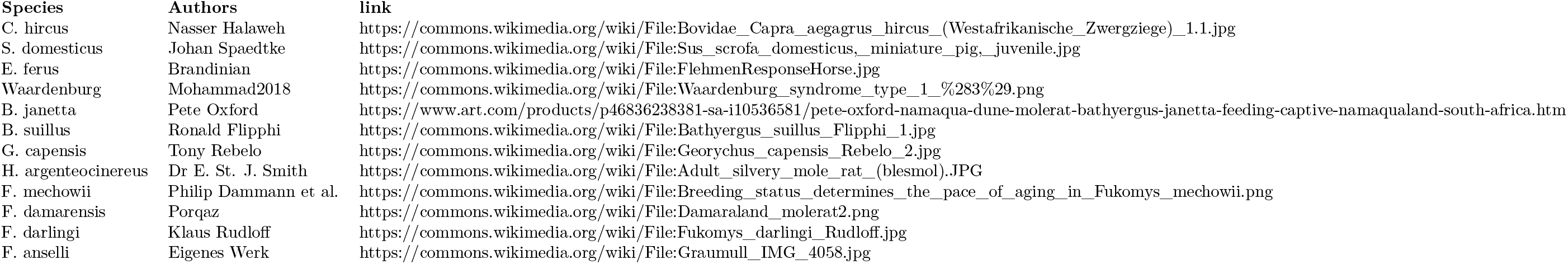
Authorship of images individual images in Fig. 2.

